# Potentiation of Acetamiprid Toxicity by Herbicide and Fungicide: Triple Mixture Outperforms Binary Combination

**DOI:** 10.1101/2025.10.15.682505

**Authors:** Jan Erik Sedlmeier, Emma Gamer, Birgit Höglinger, Frank Walker, Carsten A Brühl, Georg Petschenka

## Abstract

Pesticides are considered a major driver of insect decline, and non-target organisms in agricultural landscapes are frequently exposed to complex mixtures of agrochemicals. However, despite extensive research on mixture toxicity, most studies have predominantly focused on binary combinations, well-known synergistic interactions, and insecticide-fungicide mixtures, while other pesticide classes such as herbicides and higher-order mixtures remain underrepresented. In this study, we investigated the effects of binary and ternary combinations of representative active substances from three major pesticide classes: acetamiprid (insecticide), fluopyram (fungicide), and terbuthylazine (herbicide). We conducted acute contact dose-response assays on the non-target insect *Oncopeltus fasciatus*. While only acetamiprid showed acute toxicity when applied individually, its combination with terbuthylazine significantly increased toxicity, reducing the median effective dose (ED_50_) by a factor of 2.7. A similar, although not statistically significant, trend was observed for the combination with fluopyram. The mixture of all three compounds resulted in a marked increase in toxicity, exceeding the effects observed in any binary or single-substance treatment. These mixture effects deviate from classical additivity assumptions and can be interpreted as a potentiation of acetamiprid toxicity. While potential causes are discussed, the underlying mechanisms remain unclear. Our findings add to the growing body of evidence suggesting that non-additive effects beyond well-established pesticide combinations may occur more frequently than previously assumed. The pronounced response in the ternary mixture further indicates that higher-order mixtures may pose particular challenges for environmental risk assessment. Although this study follows a conceptual approach, the selected substances and concentrations are aligned with agricultural use patterns. Overall, our results highlight the potential for unexpected mixture effects across pesticide classes and emphasize the need to account for such interactions in environmental risk assessment.

## 1. Introduction

Pesticide contamination of the environment caused by intensified agriculture is considered one of the major drivers of the unprecedented decline in insect biomass, species richness and abundance (Geiger et al. 2010; Sánchez-Bayo and Wyckhuys 2019; Wagner et al. 2021; Dicks et al. 2021). While the ecological effects of individual active substances have been widely studied, non-target organisms in agricultural landscapes are rarely exposed to single compounds alone. Instead, they are typically confronted with complex mixtures of agrochemicals originating from formulated products, tank mixtures, or successive applications of different pesticides.

Recent studies increasingly document a widespread and year-round contamination of landscapes with such mixtures across multiple environmental matrices, including soil, vegetation, water and air (e.g. Kruse-Plaß et al. 2021; Silva et al. 2023; Brühl et al. 2024; Mauser et al. 2025). These mixtures often comprise combinations of active substances from different pesticide classes, such as insecticides, fungicides and herbicides, and are detected not only in agricultural areas but also in protected habitats. Although concentrations of individual compounds are often low, their co-occurrence can result in considerable cumulative exposure levels. From an ecological perspective, this raises concerns about so-called cocktail effects in non-target organisms (Belden et al. 2007; Cedergreen 2014; Martin et al. 2021).

Mixture toxicity is commonly predicted based on toxicity data of individual substances using established concepts such as concentration addition (CA). This approach assumes that chemicals contribute additively to the overall toxicity and is generally considered a conservative and sufficiently accurate model, even for compounds with different modes of action (Belden et al. 2007; Cedergreen 2014). Accordingly, many experimentally observed mixture effects follow additive expectations. Despite this, deviations from additivity can occur, resulting in synergistic or antagonistic interactions due to specific biochemical or physiological interactions between compounds (Cedergreen 2014). However, synergistic effects are typically associated with specific compound groups, and their underlying mechanisms are well understood. Notably, cholinesterase inhibitors and azole fungicides are involved in approximately 95% of reported synergistic interactions in binary pesticide mixtures (Cedergreen 2014).

Although such non-additive effects are comparatively rare, their occurrence challenges the predictive power of current risk assessment approaches, particularly as mixture complexity increases (Backhaus and Faust 2012). At the same time, despite the substantial body of research on mixture toxicity (Belden et al. 2007; Cedergreen 2014; Martin et al. 2021), existing studies are subject to several biases. Research has predominantly focused on aquatic systems and, in terrestrial contexts, on pollinators such as bees (Tosi et al. 2022). In addition, mixture studies are often restricted to binary combinations and tend to focus on well-known synergistic interactions, particularly involving insecticides and fungicides (Belden et al. 2007; Tosi et al. 2022). In contrast, herbicides, despite their widespread use and frequent detection in agricultural landscapes, have received comparatively little attention in the context of mixture toxicity (Tosi et al. 2022).

In this study, we address this research gap by investigating combined effects of active substances representing the three major pesticide classes: insecticides, fungicides and herbicides. Rather than aiming to replicate a specific environmental exposure scenario, we followed a conceptual approach using representative surrogate compounds. At the same time, the selected substances and tested concentrations were chosen to aligned with agricultural practices, thus enhancing the relevance of the findings to real-world conditions.

As surrogates, we selected the insecticide acetamiprid (CAS: 135410-20-7), the fungicide fluopyram (CAS: 658066-35-4) and the herbicide terbuthylazine (CAS: 5915-41-3). Acetamiprid is a neurotoxic neonicotinoid insecticide that can act as a contact or systemic insecticide. It binds to the post-synaptic nicotinic acetylcholine receptor, disrupting neural transmission (IRAC Class 4A; IRAC n.d.). It is currently the only compound of this class still approved for outdoor use in the European Union. Despite being considered less harmful than previously banned neonicotinoids, concerns remain regarding its effects on non-target insects (Varga-Szilay and Tóth 2022; Sedlmeier et al. 2025). Fluopyram is a succinate dehydrogenase inhibitor (SDHI; FRAC Class C2) that disrupts mitochondrial respiration by inhibiting complex II of the electron transport chain (Sun et al. 2005; FRAC 2025), while terbuthylazine is an s-triazine herbicide (HRAC Class C1) that interferes with photosystem II, leading to oxidative damage in plant cells (HRAC 2025). These compounds represent widely used pesticides in European agriculture and are frequently detected in environmental monitoring studies. In particular, fluopyram and terbuthylazine have been reported among the most commonly detected fungicides and herbicides in agricultural landscapes, often at comparatively high concentrations (e.g. Brühl et al. 2024; Honert et al. 2025; Mauser et al. 2025). Although these substances are not typically applied simultaneously, their co-occurrence in environmental samples, including binary and more complex mixtures, has been documented, albeit often at low concentrations. The relatively long environmental persistence of fluopyram and terbuthylazine (DT_90_ field: fluopyram 833 days; terbuthylazine 72.6 days; University of Hertfordshire 2026) further increases the likelihood of temporal overlap with other compounds such as acetamiprid, making combined exposure scenarios plausible.

To investigate the individual and combined effects of these substances, we used the large milkweed bug *Oncopeltus fasciatus* (Dallas, 1852) (Hemiptera: Heteroptera: Lygaeinae) as a model for a herbivorous non-target insect. This species can be readily reared under laboratory conditions, allowing for standardized dose–response experiments with large sample sizes. *O. fasciatus* is closely associated with milkweed (*Asclepias*, Apocynaceae), on whose seeds it feeds to sequester cardiac glycosides (Duffey et al. 1978). By testing binary and ternary combinations of the surrogate pesticides, we aim to assess whether mixtures across major pesticide classes can lead to non-additive toxic effects. This approach allows us to explore general principles of mixture toxicity beyond well-studied compound combinations and to contribute to a more comprehensive understanding of potential risks posed by pesticide mixtures in agricultural landscapes.

## 2. Methods

### 2.1 Bug rearing

*O. fasciatus* were reared in a climate chamber (APT.line™ KBF-ICH, BINDER GmbH, Tuttlingen, Germany) under long day conditions (16/8h light/dark cycle) at 27 °C and 60%rH. Nymphs were kept in 5.8 l plastic boxes covered with an insect mesh. The bottom of the box was covered with kitchen paper. Tap water was provided in a 15 ml centrifuge tube with a cotton plug in the lid and organic sunflower seeds were provided ad libitum as food. A piece of cotton wool was provided for egg laying.

### 2.2 Active substances

Analytical grade standards of acetamiprid, fluopyram and terbuthylazine were obtained commercially (acetamiprid: HPC Standards GmbH, Borsdorf, Germany; fluopyram/terbuthylazine: Merck KGaA, Darmstadt, Germany). The compounds were always stored in the dark at 4 °C in their original packaging.

The effect of the three active substances on *O. fasciatus* nymphs was first tested individually (see *Individual substance applications*). In parallel, all possible binary mixtures of the substances and the triple mixture were tested (*Mixed substance applications*).

### 2.3 Individual substance applications

For each active substance (a.s.), a permitted field concentration of a.s. derived from approved formulated products in the Germany (as of May 2023, see Supplementary Information Table 1) was used as the highest concentration in the dose-response assays. In crop protection, the amount of permitted plant protection product is fixed to a specific rate per hectare, but the water volume used for dilution can vary within a specified range (e.g. 200 to 400 liters per hectare). For fluopyram and terbuthylazine, the lowest volume of water has been used in the calculation to imitate the highest permitted amount of a.s./volume. However, the upper volume limit was used for acetamiprid because acetone was to be used as the common solvent in all assays and acetamiprid has a low reported solubility in acetone (~ 0.16 g/l).

For each a.s., a stock solution equivalent to the permitted field concentration was prepared. Fluopyram and terbuthylazine were dissolved in pure acetone, while acetamiprid was dissolved in a 9:1 acetone:deionized water mixture. The resulting stock concentrations were 83.2 ng/µl (acetamiprid), 990.0 ng/µl (fluopyram) and 3694.0 ng/µl (terbuthylazine). Detailed information on the calculations and volumes used is provided in Supplement Information Tables 2-4. Stock solutions were stored in transparent glass vials sealed with metal screw caps and kept in the dark at 4 °C. A dose-range series of six concentrations was prepared for each a.s., beginning with the stock solution and diluting each subsequent concentration by a factor of three (see table 1). In addition, solvent controls were prepared for each a.s. (fluopyram/terbuthylazine: pure acetone; acetamiprid: 9:1 acetone:deionized water). Dilutions were made by pipetting 250 µl of the respective stock into 1.5 ml ND8 amber glass screw-neck vials (LABSOLUTE—TH GEYER) pre-filled with 500 µl of pure acetone. This step was repeated to generate the full dose-range series, which was prepared once per substance and used for all individual substance assays. Each vial was thoroughly vortexed immediately after every dilution step.

**Table 1.** Dilution series of acetamiprid, fluopyram and terbuthylazine used in the *O. fasciatus* individual substance dose-response assays. Each concentration is equivalent to the dose applied topically to *O. fasciatus* nymphs. The highest dose corresponds to the permitted field concentrations of terbuthylazine and fluopyram, based on approved formulated products in Germany (as of May 2023). For acetamiprid, the field-use concentration could not be reached because of its lower solubility in acetone compared to water, resulting in a lower applied concentration.

| Field<br>Concentration<br>Equivalent [%] | Active Substance [ng/μl] |  |  |
| --- | --- | --- | --- |
|  | Acetamiprid | Fluopyram | Terbuthylazine |
| 100 | 83.2 | 990.0 | 3694.0 |
| 33.3 | 27.7 | 329.7 | 1230.1 |
| 11.1 | 9.2 | 109.9 | 410.0 |
| 3.7 | 3.1 | 36.6 | 136.7 |
| 1.24 | 1.0 | 12.3 | 45.8 |
| 0.41 | 0.3 | 4.1 | 15.1 |
| 0.0 | 0.0 | 0.0 | 0.0 |

For each dose-response assay, L4 nymphs (i.e. 4th instar nymphs) of *O. fasciatus* were isolated from the rearing box and transferred to Petri dishes. Six nymphs were randomly assigned to one of the six treatments (i.e. concentrations) and placed in individual 90 mm Petri dishes. Immediately before topical pesticide application, the nymphs were anesthetized with CO_2_. Then, 1 µl of a.s. solution (or pure solvent for the controls) was applied dorsally on the abdomen of each nymph using a Hamilton repeating dispenser with a 50 µl Hamilton syringe (Hamilton Bonaduz AG, Switzerland). All a.s. solutions were thoroughly vortexed immediately before drawing the solution into the syringe. Application started with the control followed by increasing concentrations of the a.s. solutions. The syringe was rinsed six times with pure acetone between concentrations. After the droplet had completely evaporated from the bug’s body surface and the nymphs had recovered from anesthesia, each bug was transferred to an individual 55 mm petri dish containing filter paper at the bottom, sunflower seeds and water in a 0.5 ml Eppendorf tube plugged with cotton wool. All Petri dishes within the same a.s. assay were randomized and placed in the same climate chamber under the conditions described above. The number of concentrations, individuals and cohorts tested are detailed in Table 2.

**Table 2.** Overview of *O. fasciatus* assays performed for individual active substances assays. The concentrations tested are listed in Table 1. For the acetamiprid individual assay, 12 individuals per concentration were tested in the first cohort, but only 6 in the second due to greater initial availability of test individuals. This reduced sample size was maintained for the fluopyram and terbuthylazine assays. Abbreviations: A = acetamiprid, F = fluopyram, T = terbuthylazine. The concentrations tested are listed in Table 1.

| Assay<br>(Individual) | Acetamiprid | Fluopyram | Terbuthylazine | Number of<br>Concentrations<br>Within Dose<br>Range | Individuals<br>Tested per<br>Concentration | Total<br>Number of<br>Individuals<br>Tested | Number<br>of<br>Cohorts |
| --- | --- | --- | --- | --- | --- | --- | --- |
| A | dose range | – | – | 7 | 12/6 | 126 | 2 |
| F | – | dose range | – | 7 | 6 | 84 | 2 |
| T | – | – | dose range | 7 | 6 | 84 | 2 |

### 2.4 Mixed substance applications

Dose-response assays with a.s. mixtures were conducted for the following combinations: (1) acetamiprid + fluopyram (AF), (2) acetamiprid + terbuthylazine (AT), (3) acetamiprid + fluopyram + terbuthylazine (AFT, Table 3). Each assay consisted of the six concentration levels of acetamiprid already used in the single-compound experiments, to which the highest concentration of either fluopyram (990.0 ng/µl), terbuthylazine (3694.0 ng/µl) or a combination of the two from the single-compound experiments was added. The dose-concentrations of fluopyram and terbuthylazine were selected based on their maximum permitted field application rates in Germany and were not expected to cause pronounced acute toxic effects in *O. fasciatus* in single-compound assays. This assumption is based on the low reported acute toxicity of both substances to the honeybee (LD_50_ > 102.3 µg/bee for fluopyram and > 32 µg/bee for terbuthylazine; University of Hertfordshire 2026), although direct comparability between species is limited. A solvent only control (pure acetone) was also included.

**Table 3.** Overview of *O. fasciatus* assays performed for mixed active substance assays. A “fixed dose” indicates that the corresponding maximum permitted field concentration was used. Abbreviations: A = acetamiprid, F = fluopyram, T = terbuthylazine.

| Assay (mixture) | Acetamiprid | Fluopyram | Terbuthylazine | Number of Concentrations Within Acetamiprid Dose Range | Individuals Tested per Concentration | Total Number of Individuals Tested | Number of Cohorts |
| --- | --- | --- | --- | --- | --- | --- | --- |
| AF | dose range | fixed dose | – | 7 | 6 | 126 | 3 |
| AT | dose range | – | fixed dose | 7 | 6 | 126 | 3 |
| AFT | dose range | fixed dose | fixed dose | 7 | 6 | 126 | 3 |
| FT | – | fixed dose | fixed dose | – | 6 | 12 | 1 |

To prepare the binary substance combinations (AF and AT), an acetamiprid stock solution was first made at twice the concentration used in the individual substance assay (166.4 ng/µl, i.e. 2 x the permitted field concentration), maintaining a 9:1 ratio of acetone and deionized water. A dose-range series of six concentrations, starting from and including the acetamiprid stock solution, was then prepared as described for the individual substance assays. Consequently, all six concentrations in this dilution series were initially twice as high as those required for testing. In parallel, separate stock solutions of fluopyram and terbuthylazine were prepared in pure acetone at twice their respective maximum permitted field concentrations (i.e. 1980.0 and 7388.0 ng/µl). For each dose of the binary mixtures, 50 µl of one of the six acetamiprid dilutions was pipetted into an empty 1.5 ml ND8 amber glass vial, followed by 50 µl of the fluopyram or terbuthylazine stock solution. Mixing equal volumes (2x 50 µl) yielded the same dose range of acetamiprid as was used in the single substance assay combined with the maximum permitted field concentration of either fluopyram or terbuthylazine.

For the triple combination (AFT), an acetamiprid stock solution was first prepared at three times the concentration used in the individual substance assay (249.6 ng/ul, i.e. 3 x permitted field concentration), maintaining a 9:1 acetone-to-deionized water ratio. A six-point dose-range series, starting from and including this stock solution, was then prepared as described for the individual substance assays. As a result, all concentrations in this series were initially three times higher than the final required levels. In parallel, fluopyram and terbuthylazine stock solutions were prepared in pure acetone at 3 x their maximum field concentrations (2970.0 ng/µl and 11,082.0 ng/µl, respectively). For each dose, 50 µl of each of the six acetamiprid dilutions was pipetted into an empty 1.5 ml ND8 amber glass vial, followed by 50 µl each of the fluopyram and terbuthylazine stock solutions. Mixing equal volumes (3 x 50 µl) yielded the intended acetamiprid dose range, each combined with the maximum permitted field concentration of fluopyram and terbuthylazine.

The double- and triple-concentrated acetamiprid dose-range series, along with the fluopyram and terbuthylazine stock solutions, were used for all assays and stored in the dark at 4°C. Mixtures of A + F, A + T or A + F + T were freshly prepared for each assay.

Topical application of pesticide solutions on L4 *O. fasciatus* nymphs was carried out as described for the single substance assays above. Again, Petri dishes containing L4 nymphs were randomized within the same assay and maintained in the climate chamber under the same conditions. Each concentration in the mixture assays (and a control of pure acetone) was tested on six individuals per acetamiprid concentration and each experiment was replicated three times (n = 126 in total).

In addition to the dose-range series of acetamiprid supplemented with fluopyram or terbuthylazine, a mixture containing only fluopyram and terbuthylazine (each at 100% field application rate) was tested in a supplementary experiment with reduced replication. A volume of 10 µl of each double-concentrated stock solution (200% field concentration) was pipetted into a 1.5 ml ND8 glass vial, resulting in a 100% field concentration FT mixture. Six L4 nymphs were treated with 1 µl of the mixture and six with 1 µl of pure acetone as a control. The application was conducted as described for the individual assays above. Petri dishes containing nymphs were randomized and stored in the climate chamber under the conditions described above.

The number of concentrations, individuals and cohorts tested in all mixed substance assays are detailed in Table 3.

### 2.5 Evaluation of toxic effects

Nymphs were evaluated 96 hours after application. Petri dishes were removed from the climate chamber and placed in the laboratory. Each nymph was classified into one of three categories: dead, affected or unaffected. Classification was aided by gently touching the abdomen with a fine brush. Healthy individuals typically responded with a flight reaction or flinching. Affected individuals reacted slowly, if at all, often only moving a few legs or antennae, or leaning to one side. Nymphs that appeared dead (typically lying on their back with bent legs and dark discoloration) were further tested with the brush to confirm the absence of any response. This evaluation scheme was applied consistently across all assays. Raw data for all treatments is provided in Supplementary Data.

### 2.6 Statistical analysis and graphical representation

Using R (version 4.3.0; R Core Team 2025) and R-Studio (version 2023.6.1.524; Posit team 2026) and the ‘drc’ package (Ritz et al. 2015), a four-parameter log-normal cumulative dose-response model, assuming a binomial distribution, was fitted to the response data at 96 h post application for the data from the individual acetamiprid assay and from the mixed assays AF and AT.

The following data were not included in the model: (1) data from the individual fluopyram and terbuthylazine assays as no effects were observed; (2) data from the fluopyram and terbuthylazine mixed assay (FT) as no effects were observed; (3) data from the acetamiprid, fluopyram and terbuthylazine triple mixed assay (AFT) as the effects were too strong across the dose range (see Results: Mixed substance assays). The lower asymptote of the dose-response model was set to ‘0’ because in all included assays, none of the control animals showed any effects or mortality. The upper asymptote was set to ‘1’. The *plot()* function was used to generate dose-response curves. Model fit was assessed by visual inspection of the regression curve and comparison of the Akaike Information Criterion (AIC) values. LD_50_ values (and respective 95% confidence intervals) were derived from the model using the *ED()* function. The *compParm()* function was used to assess differences in the dose required to affect 50% of the tested population (ED_50_ value) The function compares individual curves (i.e., ED_50_ values) within the model and performs approximate t-tests with α = 0.05. Note that all derived ED_50_ values, including those from the mixed assays, refer to the respective acetamiprid concentration in ng/µl. The full R-code is provided in Supplementary Information – Supplementary Code.

Figure 1, including the dose response curves and mean data points from the AFT treatment, was created in R and later edited in Inkscape (version 1.4; Inkscape Project 2025). Figure 2 was created with JMP® Student Edition (version 18.2.2; JMP Statistical Discovery LLC 2025) and later edited with Inkscape.

**Figure 1.**
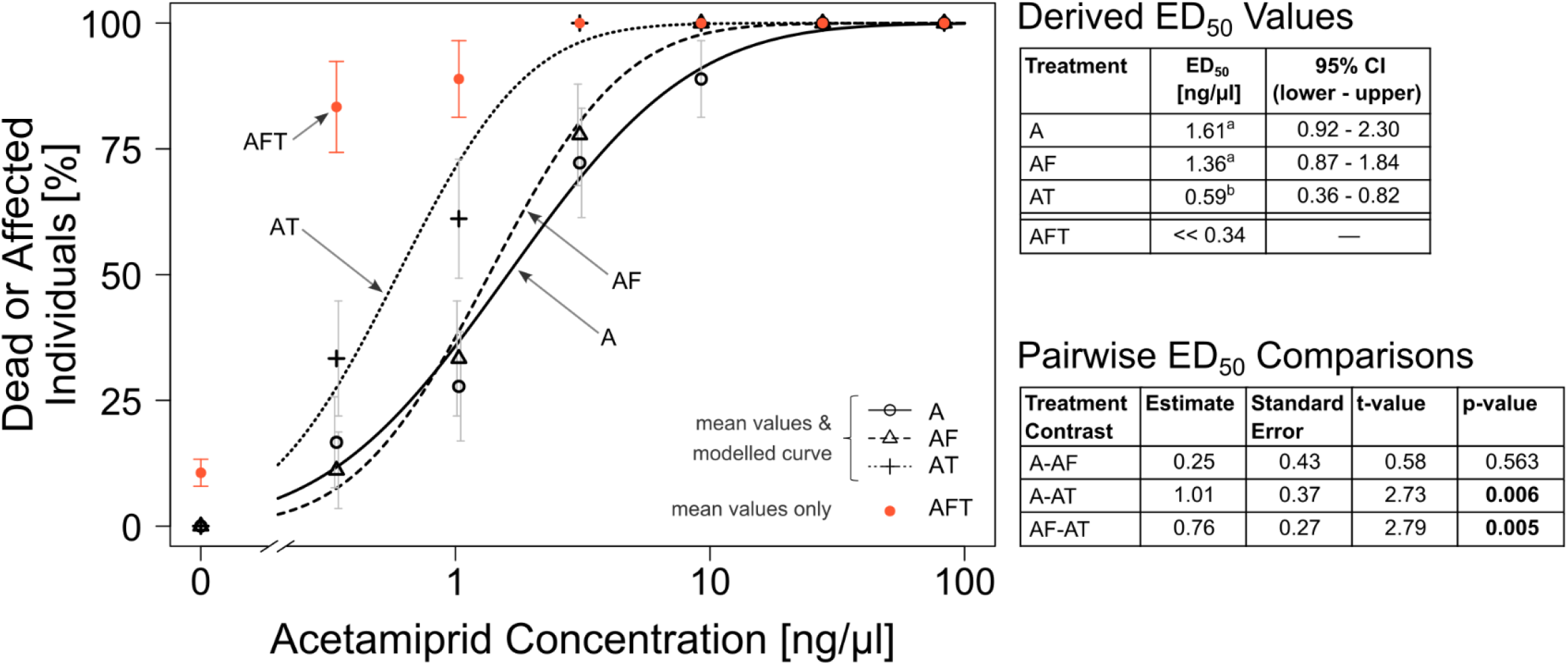
*Oncopeltus fasciatus* dose-response curves for individual and mixture assays and comparison of derived ED_50_ values. A: acetamiprid individual assay, AF: acetamiprid + fluopyram, AT: acetamiprid + terbuthylazine, AFT: acetamiprid + fluopyram + terbuthylazine. Dose-response relationships were modelled simultaneously using a four-parameter log-normal function. Black symbols represent treatment-specific mean responses per concentration, with grey error bars indicating standard errors (SE). SE indicators were offset in cases of overlap to facilitate interpretation. Red data points (and respective SE indicators) represent mean responses to the AFT treatment but were not included in the dose-response model due to convergence failure. Derived ED_50_ values and respective 95% confidence intervals (CI) for each treatment included in the model are listed in the top right table. For AFT, the ED_50_ is reported as << 0.34 because 83.3% mean response was observed already at the lowest tested concentration (0.34 ng/µl), indicating that the actual ED_50_ must lie well below this value. LD_50_ values derived from the model were compared using the *compParm* function of the *drc* package in R. LD_50_ values of treatments not sharing a common letter are significantly different. A statistical report on the pairwise comparisons is given in the bottom right table with bolt p-values indicating significant differences at α = 0.05. All numbers presented in the adjacent tables are rounded. For all assays, N = 18 biological replicates (nymphs) per concentration

**Figure 2.**
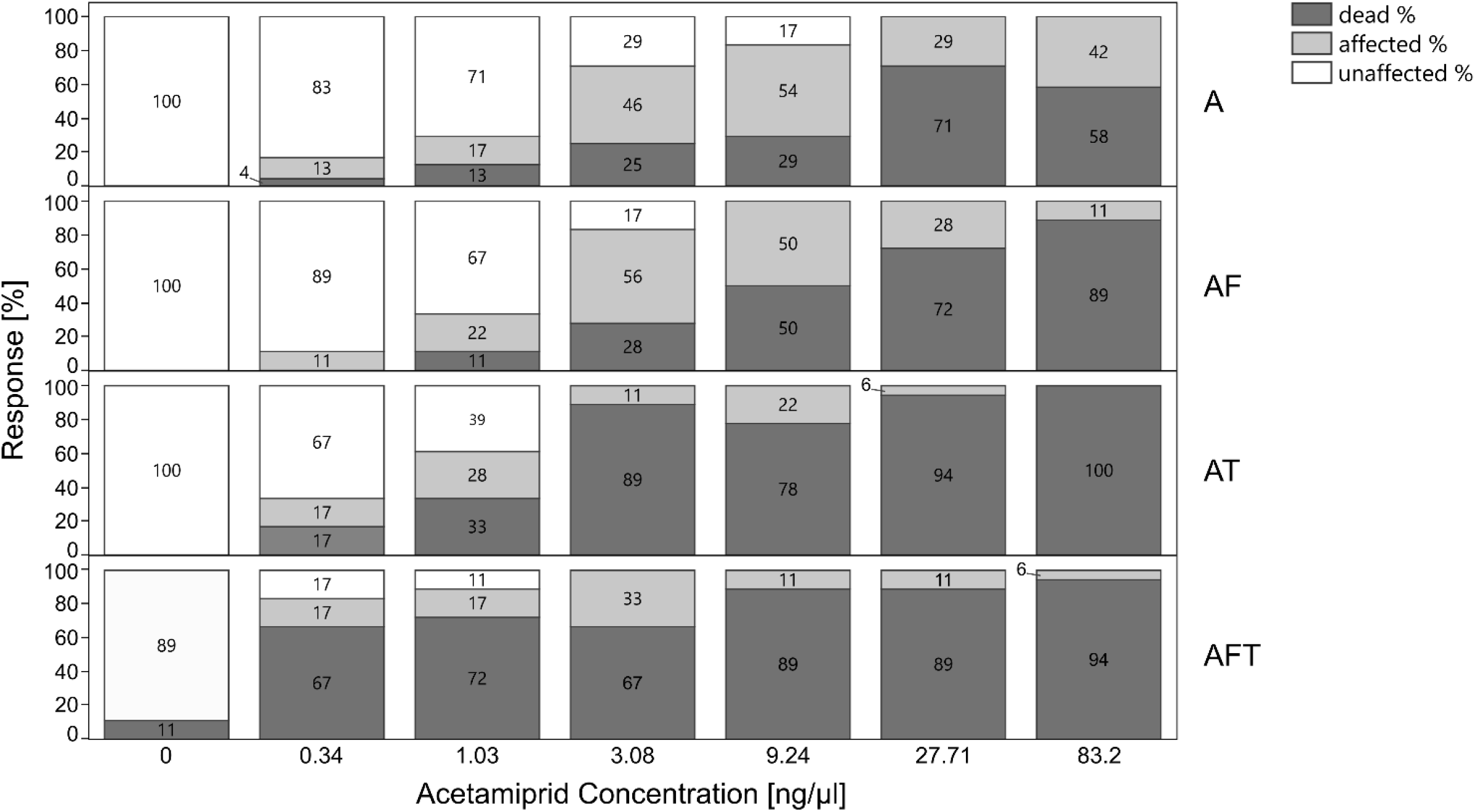
Response of *O. fasciatus* nymphs to individual and mixed substance treatments containing acetamiprid. Proportions of response types are indicated by different colors for each acetamiprid concentration tested. Numbers inside or left of the bars indicate percentage response (rounded). For all assays, N = 18 biological replicates (nymphs) per concentration

## 3 Results

### 3.1 Individual substance assays

*Oncopeltus fasciatus* nymphs showed high sensitivity to the insecticide acetamiprid 96 hours after application. At all concentrations tested, dead or affected nymphs were observed. However, at no concentration 100% mortality was observed in the two cohorts tested. The derived ED_50_ for acetamiprid is 1.61 ng/µl (combined ‘*dead or affected*’ data) based on a multiple treatment cumulative dose-response model with a log-normal distribution (Fig. 1).

No effects were observed across concentrations in the single-substance dose-response assays with fluopyram and terbuthylazine 96 hours after application, and no dose-response curves were fitted. However, there were isolated cases of mortality in both experiments. Two nymphs died in the fluopyram assay, one at 33.3% maximum field concentration equivalent (1^st^ cohort) and one at 11.1% (2^nd^cohort). In the terbuthylazine assay, four nymphs died, two in the control (one in each cohort), one at 3.7% field concentration equivalent (1^st^ cohort) and one at 1.24% (2^nd^cohort). However, in both assays, there were no nymphs categorized as affected in addition to those that died, supporting that these were random mortality events and these values were not considered further.

### 3.2 Mixed substance assays

The effect of the AF mixed treatment (insecticide + fungicide) on the nymphs compared to the individual acetamiprid application was not significantly different based on the cumulative dose-response model followed by an ED_50_ comparison (t=0.58, p=0.563; Fig. 1). However, sensitivity towards the AT mixed application (insecticide + herbicide) compared to both acetamiprid alone and the AF treatment was significantly higher based on the ED_50_ comparison (A-AT: t=2.73, p=0.06; AF-AT: t=2.79, p=0.005). The ED_50_ of AT (0.59 ng/µl) was approximately 2.7-fold lower than that of acetamiprid alone (1.61 ng/µl), and 2.3-fold lower than that of AF (1.36 ng/µl). The strongest effects were observed in the AFT treatment (insecticide + fungicide + herbicide). However, the data could not be included in the dose-response model because the observed effects were too pronounced across the entire dose range. At the lowest tested dose (0.34 ng acetamiprid /µl), the mean response was 83.3%, exceeding the inflection point of a typical dose-response curve. As a result, no curve could be fitted. Nonetheless, this suggests that the ED_50_ for the AFT treatment would be well below 0.34 ng/µl.

When differentiating between dead and merely affected individuals, mortality in the AFT and AT assay was proportionately higher at lower acetamiprid doses than in the other treatments (Fig. 2). Strikingly, in the AFT treatment, even at the lowest concentration, 67% of nymphs died within 96 hours, compared to 17% in the AT, 0% in the AF, and 4% in the assay with acetamiprid only.

No effects on *O. fasciatus* nymphs were observed in the combined FT treatment (fungicide + herbicide, each at 100% field concentration, see Supplementary Data) 96 hours after application. All nymphs were unaffected in both the treatment and the control group.

## 4 Discussion

To investigate combined toxicity across major pesticide classes, we assessed the insecticide acetamiprid, the herbicide terbuthylazine, and the fungicide fluopyram in the non-target herbivore *O. fasciatus*. Each compound was tested individually in dose-response assays, followed by binary and ternary mixtures, revealing considerable increases in acetamiprid toxicity if combined with terbuthylazine alone or together with fluopyram.

Although non-additive effects are generally rare, they arise consistently in combinations of specific pesticides based on well characterized mechanistic principles (Cedergreen 2014; Tosi et al. 2022). A prominent example is the synergism between neonicotinoids or pyrethroids and azole fungicides which interfere with insect detoxification pathways (Sgolastra et al. 2017; Schuhmann et al. 2022). Our findings support existing indications that non-additive interactions may not be limited to specific pesticide groups such as azole fungicides (Manzer et al. 2024). Rather, they could emerge more broadly, including from combinations involving herbicides, a pesticide class less commonly associated with such effects (Tosi et al. 2022). In addition, complex mixtures (i.e., those containing three or more compounds) appear to be especially problematic due to their unpredictable interaction potential.

This is concerning because the absence of acute toxicity in single-substance applications of terbuthylazine and fluopyram would not have enabled an accurate prediction of combined toxicity based on the concept of concentration addition. Since no measurable acute effects were observed for terbuthylazine and fluopyram individually (and thus no ED_50_ was determined), the additive prediction would mirror the response of acetamiprid alone. The marked reduction in ED_50_ observed for acetamiprid in the binary mixture (A: 1.61 ng/µl; AT: 0.59 ng/µl), as well as the pronounced effects in the ternary combination (AFT), indicate a clear deviation from additivity. Because formal synergy modelling was not feasible due to the lack of measurable single-substance effects and a complete dose–response matrix, the observed interaction is best interpreted as a potentiation of acetamiprid toxicity.

In the following sections we address potential mechanisms by which terbuthylazine and fluopyram might facilitate potentiation of acetamiprid toxicity. After that, we provide context on the ecotoxicological relevance of our data while outlining limitations of this study.

### 4.1 Terbuthylazine

Herbicides have been shown to exert direct detrimental effects on non-target organisms, including invertebrates (Wan et al. 2025). However, the mode of action of terbuthylazine does not immediately suggest off-target effects in insects, as it specifically targets the photosystem II in plants. While adverse effects of terbuthylazine have been reported in both vertebrates and invertebrates, including impacts on antioxidant defense systems and the induction of oxidative stress (Semren et al. 2018), comparable data for insects are currently lacking. While such effects may become relevant under higher exposure or in mixture scenarios, the lack of acute toxicity our single-compound assays limits further interpretation in this context.

At present, there is no well-established mechanistic framework to explain the observed mixture effects. S-triazines such as terbuthylazine are known to exert synergistic effects in combination with specific organophosphate insecticides that require cytochrome P450-mediated bioactivation to their more potent oxon forms (Pérez et al. 2013; Cedergreen 2014). These interactions are attributed to the ability of s-triazines to induce P450 enzymes in non-target organisms, despite their primary mode of action targeting photosynthesis in plants. However, the direction of these effects is not consistent across compounds and species, as both synergistic and antagonistic interactions have been reported (Kao et al. 1995). In contrast, acetamiprid does not require bioactivation to exert its toxicity, and therefore the classical mechanism described for organophosphates cannot be applied in this study. Nevertheless, the ability of s-triazines to modulate P450 activity suggests that interactions with detoxification pathways may still occur. However, while inhibition of P450-mediated detoxification would represent a plausible mechanism for the increased toxicity of acetamiprid (Iwasa et al. 2004), there is currently no evidence that terbuthylazine exerts such effects in *O. fasciatus* or other insect species.

An alternative explanation is that co-exposure to terbuthylazine imposes an additional xenobiotic burden, thereby reducing the overall capacity of detoxification systems. In our experiments, 3.7 µg terbuthylazine was co-applied at each acetamiprid concentration, representing a substantial additional chemical load for the nymphs. At an average body weight of approximately 50 mg per L4 nymph, this corresponds to a nominal exposure of approximately 0.07 µg per mg body weight, indicating a considerable additional chemical load. The reported acute contact LD_50_ of terbuthylazine in honey bees (> 32 µg/bee; University of Hertfordshire 2026) indicates low intrinsic toxicity, which is consistent with the absence of acute effects observed in *O. fasciatus* following exposure to terbuthylazine alone. However, even in the absence of direct toxicity, such additional exposure may still interfere with physiological processes involved in detoxification.

In summary, the mechanism underlying the observed potentiation of acetamiprid toxicity remains unclear. Both interference with detoxification pathways and an increased overall xenobiotic burden represent plausible explanations, but these hypotheses require further experimental validation.

### 4.2 Fluopyram

Fluopyram did not cause acute toxicity in *O. fasciatus* nymphs in our study. Likewise, the combination with acetamiprid did not result in a statistically significant increase in toxicity compared to acetamiprid alone. To our knowledge, no consistent evidence for non-additive interactions between fluopyram and insecticides has been reported. However, interactions involving mitochondrial stressors have been described for other fungicides. For example, Manzer et al. (2024) demonstrated increased acetamiprid toxicity in honeybees following larval exposure to a mixture of two fungicides targeting mitochondrial electron transport, including the SDHI boscalid. Notably, these effects were delayed and only became apparent in adult stages, suggesting that acute toxicity assays may underestimate mixture effects involving SDHI fungicides.

The succinate dehydrogenase is a key enzyme of the mitochondrial respiratory chain involved in ATP production. As this enzyme is conserved across eukaryotes, its inhibition by fungicides such as fluopyram would not be limited to fungi. This assumption is supported by its use as a nematicide with the same mode of action (Schleker et al. 2022), and by findings showing that the SDH of the honeybee Apis mellifera is sensitive to fluopyram (Bénit et al. 2019). Inhibition of mitochondrial respiration could theoretically reduce cellular energy availability, which is essential for detoxification processes such as biotransformation and active transport of xenobiotics. In addition, exposure to fluopyram has been associated with oxidative stress and reduced fitness in Caenorhabditis elegans (Liu et al. 2022), as well as metabolic disruptions in other taxa, indicating broader cellular stress responses. Such effects may not directly manifest as acute toxicity in single-substance assays but could increase susceptibility to insecticides. In line with this, a steeper decline in the proportion of unaffected nymphs with increasing acetamiprid concentrations was observed in the mixture treatment compared to acetamiprid alone. While this observation does not provide conclusive evidence for non-additive interaction, it may indicate subtle effects not captured by ED_50_-based analysis.

In summary, the mode of action of fluopyram and its potential to induce metabolic stress suggest that it may contribute to increased toxicity under more complex mixture conditions.

### 4.3 Complex mixtures

Even at the lowest acetamiprid concentration, effect levels observed in in the triple mixture (AFT) exceeded the ED_50_ observed when acetamiprid was applied alone. While the mechanisms behind this potentiation remain unknown, a plausible explanation lies in the simultaneous disruption of multiple physiological systems. As established above, terbuthylazine may contribute to an increased xenobiotic burden and potentially interfere with detoxification pathways. At the same time, fluopyram is known to impair mitochondrial respiration and thus cellular energy production (Bénit et al. 2019). In combination, these effects could reduce the capacity of detoxification systems, which rely on ATP-dependent processes such as biotransformation and active transport of xenobiotics. If detoxification were already burdened by other xenobiotics, potentially impaired, and ATP production were additionally disrupted, even low concentrations of acetamiprid may result in disproportionately strong neurotoxic effects. The observed response in the AFT treatment is therefore consistent with a scenario in which multiple stressors converge, leading to a breakdown of physiological homeostasis not observed in binary or single applications.

Importantly, neither fluopyram nor terbuthylazine alone, nor their respective binary combinations with acetamiprid, resulted in comparable increases in toxicity. This indicates that the observed effect is not driven by a single dominant interaction, but rather emerges from the combined action of all three compounds. The results indicate that the toxicity of the triple mixture cannot be adequately explained by simple additive models, supporting previous suggestions that especially higher-order mixtures may exert effects that are not reliably predictable on existing single substance data (Backhaus and Faust 2012). This highlights the risks posed by complex pesticide mixtures as well as the limitations of current risk assessment approaches, which still mainly focus on single substance toxicity along with the assessment of combination products, common tank mixtures or known synergistic interactions (e.g. pyrethroids & piperonyl butoxide, pyrethroids or neonicotinoids & azole fungicides; (Backhaus and Faust 2012; EFSA PPR 2015).

It should be noted that the present study focused on acute toxicity. Sublethal effects, including behavioral impairment, developmental disturbances, or reproductive deficits, were not assessed but are frequently reported for pesticide mixtures (Müller 2018; Gandara et al. 2024). Given the strong acute response observed in the AFT treatment, it is conceivable that such effects may be even more pronounced under chronic or environmentally realistic exposure scenarios. This underscores the need for further studies addressing complex mixture effects across different endpoints and life stages.

### 4.4 Limitations and Realistic Scenarios

Several limitations of the present study should be considered. The experiments focused on acute toxicity in a single species under controlled laboratory conditions, and did not address chronic, sublethal, or life-stage-specific effects. In addition, the tested exposure scenarios represent simplified combinations of up to three compounds at concentrations relevant to field application, whereas environmental mixtures are typically more complex and variable. Despite these limitations and the fact that this study was not designed to reflect a specific local exposure scenario, but rather to investigate general patterns of combined toxicity across pesticide classes, the mixtures tested here are not merely hypothetical. There are realistic conditions under which non-target organisms may encounter comparable combinations of active substances:

1. Non-target organisms may encounter low concentrations of one compound (e.g. acetamiprid), for example due to off-field transport or degradation after application, and subsequently be exposed to higher concentrations of another compound through a new application event. Such a scenario would mirror the AFT mixture, where even at low acetamiprid concentrations, the combination with terbuthylazine and fluopyram at field-realistic concentrations significantly elevated acute toxicity.
2. While environmental concentrations of individual pesticides are often low, the exposure of non-target insects to complex mixtures including different modes of action are a common feature of agricultural landscapes rather than the exception (Brühl et al. 2021). Recent environmental monitoring has detected a.s. combinations corresponding to those tested in our experiments in agricultural landscapes. For example, the binary AT mixture which showed a potentiated acetamiprid toxicity, was detected in topsoil samples from agricultural sites in Rhineland-Palatinate (Germany) (Honert et al. 2025). While the full combination of all three substances (AFT) was detected less frequently, fluopyram and terbuthylazine were commonly found alongside other neonicotinoids, indicating that similar multi-component exposure scenarios are likely to occur. Although these are environmental residues that do not compare to the concentrations tested in this study, our results suggest an ecological risk posed by this specific combination. In addition, we only tested acute toxic effects. Mixtures of lower concentrations might, result in sublethal effects (Bartling et al. 2024; Gandara et al. 2024).

There is a growing body of evidence that unpredictable interactions of pesticides may emerge predominantly in complex mixtures and from combinations previously not associated with such effects (Gandara et al. 2024; Manzer et al. 2024). This represents a challenging aspect of environmental risk assessment. At the same time, it is important to recognize that the systematic evaluation of all possible mixture combinations is not feasible, given the large number of compounds and the variability of environmental exposure scenarios. While mixture toxicity can often be predicted with reasonable accuracy using established concepts, experimental testing of all conceivable combinations would be impractical (Kortenkamp et al. 2009). This highlights an inherent tension between the complexity of real-world exposure and the need for robust and feasible risk assessment approaches. However, our findings underline the importance of considering unexpected interactions when assessing environmental risks posed by pesticides.

## Acknowledgments

We would like to thank Sarah Rißmann for her consistent support throughout the laboratory work and insect rearing. We would also like to thank the Graduate Academy at the University of Hohenheim for making this project possible by awarding a scholarship as part of the Baden-Württemberg Graduate Support Program to JES.

## Data Availability

All data and statistical code generated in this study are provided in the supplementary materials.

## Statements and Declarations

### Contributions

J.E.S., E.G. and G.P designed the experiments. E.G. and J.E.S. conducted the experiments. B.H. and F.W. developed the active substance mixing strategies in a common solvent based on chemical properties. J.E.S. and E.G. performed the analysis of the experiments. J.E.S. wrote the original draft of the manuscript and created the figures. G.P. supervised the research and edited the manuscript.

