## Supplementary Information for "Potentiation of Acetamiprid Toxicity by Herbicide and Fungicide: Triple Mixture Outperforms Binary Combination"

**This document includes:**

### Supplemantary Table 1: Derivation of the field concentrations used for the three active ingredients based on products approved in Germany as of May 2023.

| **Active Ingredient** | **Product Name** | **Registration Number (BVL Germany)** | **Active Ingredients Contained** | **Field Application Rate** | **Crop** | **Application Number (BVL Germany)** | **Equivalent Field Concentration Used** |
| --- | --- | --- | --- | --- | --- | --- | --- |
| acetamiprid | Mospilan® SG | 005655-00 | 200 g/kg acetamiprid | 250 g/ha in 200-600 l/ha | potato | #005655-00/06-002 | 83.2 ng/µl |
| fluopyram | Silvron® | 00A457-00 | 100 g/l fluopyram 100 g/l bixafen | 1 l/ha in 100-400 l/ha | spelt | #00A457-00/00-011 #00A457-00/00-012 | 990 ng/µl |
| terbuthylazine | Spectrum® Gold | 006380-00 | 250 g/l terbuthylazine 280 g/l dimethenamide-P | 3 l/ha in 200-400 l/ha | corn | #006380-00/00-001 | 3694 ng/µl |

### **Supplementary Table 2:** Preparation of active substance stock solutions – acetamiprid.

|  | **Acetamiprid** | | | | | |
| --- | --- | --- | --- | --- | --- | --- |
|  | **target concentration [mg/ml]** | **target volume [ml]** | **weigh-in [mg]** | **calculation of total solvent volume [ml]** | **water [ml]** | **acetone [ml]** |
| **Individual Substance Application** | 0.0832 | 10 | 0.84 | 0.84/0.0832 = 10.096 | 1 | 9.096 |
| **Binary Mixtures (AF, AT)** | 0.1664 | 10 | 1.6 | 1.6/0.1664 = 9.615 | 1 | 8.615 |
| **Ternary Mixture (AFT)** | 0.2496 | 10 | 2.44 | 2.44/0.2496 = 9.776 | 1 | 8.776 |

### **Supplementary Table 3:** Preparation of active substance stock solutions – fluopyram.

|  | **Fluopyram** | | | | | |
| --- | --- | --- | --- | --- | --- | --- |
|  | **target concentration [mg/ml]** | **target volume [ml]** | **weigh-in [mg]** | **calculation of total solvent volume [ml]** | **water [ml]** | **acetone [ml]** |
| **Individual Substance Application** | 0.99 | 3 | 2.92 | 2.92/0.99 = 2.949 | – | 2.949 |
| **Binary Mixtures (AF, FT)** | 1.98 | 3 | 5.88 | 5.88/1.98 = 2.969 | – | 2.969 |
| **Ternary Mixture (AFT)** | 2.97 | 3 | 8.74 | 8.74/2.97 = 2.943 | – | 2.943 |

### **Supplementary Table 4:** Preparation of active substance stock solutions – terbuthylazine.

|  | **Terbuthylazine** | | | | | |
| --- | --- | --- | --- | --- | --- | --- |
|  | **target concentration [mg/ml]** | **target volume [ml]** | **weigh-in [mg]** | **calculation of total solvent volume [ml]** | **water [ml]** | **acetone [ml]** |
| **Individual Substance Application** | 3.694 | 3 | 11.05 | 11.05/3.694 = 2.991 | – | 2.991 |
| **Binary Mixtures (AT, FT)** | 7.388 | 1 | 7.79 | 7.79/7.388 = 1.054 | – | 1.054 |
| **Ternary Mixture (AFT)** | 11.082 | 3 | 32.19 | 32.19/11.082 = 2.905 | – | 2.905 |

### **Supplementary Code:** R-Script – dose-response model and creation of figure 1

library(readxl)

library(drc)

library(MASS)

dfm <- read_excel("Data_Oncopeltus.xlsx")

str(dfm)

dfm$concentration=as.numeric(dfm$concentration)

dfm$treatment=as.factor(dfm$treatment)

str(dfm)

dfm4 <- subset(dfm, time_days == 4 & !(treatment %in% c('AFT', 'F', 'T'))) # filter data

View(dfm4)

#################################################

\##	model for *dead + affected* - 4 days after application	##

#################################################

m1 <- drm(dead_affected/total ~concentration, treatment, data = dfm4, weights=total, type="binomial",

fct = LN.4(fixed = c(NA,0,1,NA), names = c("slope","lower","upper","ED50")),

pmodels = data.frame(treatment,treatment,treatment,treatment),

control=drmc(errorm=FALSE))

\# analysis of model fit

summary(m1)

plot(m1)

AIC(m1)

\# ED50 extraction and comparison

summary(m1)

ED(m1, c(50), interval = "delta")

compParm(m1, "ED50" , "-" )

#################

\##	create graph	##

#################

\## plot cumulative dose-response graph including SE bars

\# Calculate mean and standard error for each concentration and species

library(dplyr)

mean_se_data <- dfm4 %>%

group_by(concentration, treatment) %>%

summarise(

mean_response = mean(dead_affected/total),

se_response = sd(dead_affected/total) / sqrt(n())

) %>%

ungroup()

\# plot

plot(m1, broken = TRUE, xlim = c(0, 100), ylim = c(0, 1),

yt = c('0.0', 0.25, 0.5, 0.75, '1.0'),

ytlab = c('0', '25', '50', '75', '100'),

xlab = "Acetamiprid Concentration [ng/µl]",

ylab = "Dead or Affected [%]",

cex = 1,

cex.axis = 1.6,

cex.lab = 1.8,

col = 1,

lwd = 2,

legend = TRUE,

legendPos = c(90, 0.25),

type = "average")

\# Overlay mean data points & SE bars

for (spec in unique(mean_se_data$treatment)) {

\# Filter the mean and SE data for the current species

treatment_data <- mean_se_data[mean_se_data$treatment == spec, ]

\# Add error bars using the arrows function for each concentration-species pair

arrows(

x0 = treatment_data$concentration,

y0 = treatment_data$mean_response - treatment_data$se_response, # Lower end of the error bar

x1 = treatment_data$concentration,

y1 = treatment_data$mean_response + treatment_data$se_response, # Upper end of the error bar

angle = 90, code = 3, length = 0.025, col = "grey" # Customizing arrow heads and color

)

}

\# Include AFT mean values for dead_affected in same plot:

library(dplyr)

\# Calculate means for AFT*

aft_data <- dfm %>%

filter(treatment == "AFT" & time_days == 4) %>%

group_by(concentration) %>%

summarise(

mean_response = mean(dead_affected / total),

se_response = sd(dead_affected / total) / sqrt(n())

)

print(aft_data)

\# Add AFT means to plot:

points(aft_data$concentration, aft_data$mean_response,

pch = 16, col = "#FF5733")

\# Add SE indicators for AFT mean values:

arrows(

x0 = aft_data$concentration,

y0 = aft_data$mean_response - aft_data$se_response,

x1 = aft_data$concentration,

y1 = aft_data$mean_response + aft_data$se_response,

angle = 90, code = 3, length = 0.05, col = "#FF5733"

)

\## Extract mean response value for AFT at X=0 (+ SE) and plot it at X=1. Then use external program to shift it to X=0.

\# --> drc plot does not support subsequent plotting of values at x=0 outside of the own plot() function

View(aft_data) #extract y-value for x=0

points(x = 1, y = 0.1060606, pch = 16, col = "#FF5733")

\# Add SE bars

arrows(x0 = 1, y0 = 0.1060606 - 0.026902674,

x1 = 1, y1 = 0.1060606 + 0.026902674,

angle = 90, code = 3, length = 0.05, col = "#FF5733")
